# FAK inhibition suppresses breast cancer progression via DNA methylation-mediated DAB2 gene reactivation

**DOI:** 10.1101/2024.11.23.624992

**Authors:** James M. Murphy, Kyuho Jeong, Eun-Young Erin Ahn, Ssang-Taek Steve Lim

## Abstract

Epigenetic silencing of tumor suppressor genes is one of the main drivers of tumor progression. Without these tumor suppressors to reduce proliferation, tumor cells proliferate unchecked. Focal adhesion kinase (FAK) is a tyrosine kinase which is often upregulated in various tumors and promotes cell proliferation and migration. Recent studies have demonstrated that pharmacological or genetic FAK inhibition can reduce suppressive DNA methylation in vascular cells. Mechanistically, this is through nuclear FAK-mediated ubiquitination and proteasomal degradation of DNA methyltransferase 3A (DNMT3A). Treatment of breast cancer cell lines with FAK inhibitor (FAK-I) was able to reduce both FAK activity and DNMT3A protein expression. Further, global DNA methylation was reduced in breast cancer cell lines treated with FAK-I. This decrease in DNA methylation was correlated with decreased cell proliferation. We further showed that FAK-I reduced DNMT3A expression in breast cancer cells and that treatment with the proteasome inhibitor MG132 prevented loss of DNTM3A protein stability. To identify how FAK-I and DNMT3A loss could reduce breast cancer cell growth we compared RNA sequencing data from breast cancer cells treated with or without FAK-I or in shRNA DNMT3A knockdown. We have identified a potential tumor suppressor, DAB2, as being regulated by the nuclear FAK-DNMT3A axis. DAB2 is often downregulated in cancers and has been shown to play a vital role in switching TGFβ signaling from proliferative to apoptotic by altering TGFβRI binding partners. Immunoblotting and immunostaining indeed revealed that FAK-I and shDNMT3A could induce DAB2 protein expression. Further, FAK-I treatment showed efficacy in reducing tumor growth in vivo using the murine 4T1 tumor model. Immunostaining of 4T1 tumors showed FAK-I decreased DNMT3A, DNA methylation (5-methylcytosine, 5-mC), and increased DAB2 expression. Taken together, these data suggest that nuclear FAK-mediated regulation of DNMT3A can alter the epigenetic landscape and induce tumor suppressor gene expression.

## Introduction

Breast cancer is the most frequently diagnosed cancer and the leading cause of cancer death among women, accounting for 23% of the total cancer cases and 14% of the cancer deaths.^1^ One of the important classifications of breast tumors is based on the presence or absence of the estrogen receptor (ER). While most breast cancers are ER-positive, approximately 25–30% are ER-negative.^2,3^ Triple-negative breast cancer (TNBC), characterized by absence or minimal expression of ER and progesterone receptor and lack of the human epidermal growth factor receptor 2 (her2) amplification, represents 15%–20% of all breast cancers. TNBC is generally more aggressive; although many patients respond well to standard anthracycline- and taxane-based chemotherapy, long-term patient survival is poor due to high rates of relapse and recurrence.^4,5^ Therefore, there need to identify additional drugs and drug targets for TNBC patients.

Epigenetic alterations, including altered DNA methylation, are frequently detectable in human breast cancers, e.g. DNA hypermethylation mediated transcriptional silencing of tumor suppressor genes, which promotes to propagation of breast cancer cells.^6,7^ TNBC tumors show extensive promoter hypermethylation of epigenetic biomarker genes compared with other breast cancer subtypes in which promoter hypermethylation events were less frequent,^8,9^ suggesting that targeting the epigenetic machinery of TNBCs may have clinical benefits. DNA methylation is catalyzed by DNA methyltransferases (DNMTs), and typically occurs at the 5 position of cytosine (5-mC) in CG dinucleotides of CpG islands. The process that mediates DNA methylation in the genes has not been fully elucidated regarding tumor suppresser gene transcription. There are three DNMTs that can methylate cytosine: DNMT1, DNMT3A and DNMT3B. DNMT1 is a key enzyme in preserving DNA methylation during DNA replication. On the other hand, DNMT3A and DNMT3B are *de novo* DNA methylation enzymes that regulate gene expression.^10^

Although the expression of many tumor suppressors are repressed in advanced carcinoma, we identified human disabled-2 (DAB2) significantly downregulated in breast cancers and FAK-I reactivated DAB2 expression through reduced DNA methylation on *DAB2* promoter. In this study, we hypothesized that nuclear FAK could repress breast cancer proliferation and tumor growth in vivo via reducing DNMT3A stability, thereby decreasing global DNA methylation.

## Materials and Methods

### Animal experiments

Animal experiments were approved and performed according to the guidelines of the University of South Alabama and University of Alabama at Birmingham Institutional Animal Care and Use Committees. As most breast cancer cases occur in women, only female mice were used for 4T1 syngeneic experiments. Balb/c female mice were anesthetized with ketamine/xylazine and the hair was removed from around the 4^th^ inguinal mammary fat pad. The injection site was cleaned using propidium iodide followed by 70% ethanol wipe. Then 1×10^6^ 4T1 cells (in 100 μl PBS/Matrigel) were injected into the 4^th^ inguinal mammary fat pad for FAK-I experiments. After 12 days of tumor initiation, mice were treated with either vehicle (0.2% Tween-80 and 0.5% methylcellulose) or FAK-I (VS-6063, 35 mg/kg) twice daily via oral gavage until 21 days post tumor implantation. Tumors were measured twice a week using digital calipers. At the end of the experiment tumors were collected either frozen in for sectioning or used for RNA isolation.

### Reagents

Reagents for immunoblot and immunohistochemistry included FAK (#05-537), 5-mC (SAB2702243), 5-hmC (SAB2702268), Ubiquitin (#04-262), and glyceraldehyde 3-phosphate dehydrogenase (GAPDH) (#MAB374) antibodies (Millipore, Burlington, MA); DNMT3A (#SC365769), DNMT1 (#SC20701), and DAB2 (#SC136964) antibodies (Santa Cruz Biotech, Dallas, TX); pY397 FAK (#44-624G), and DNMT3B (#PA1-32317) antibodies (Thermo Fisher, Waltham, MA); MG132 and the small molecule FAK inhibitor VS-6063 (MedKoo, Morrisville, NC).

### Cell culture

All cells were grown in Dulbecco Modified Eagle Medium (DMEM) containing 10% fetal bovine serum (FBS), and 100 U/mL penicillin/streptomycin under 5% CO2 at 37°C.

### Immunoblot

Cells were lysed in RIPA buffer (pH 7.4) that included 4-(2-hydroxyethyl)-1-piperazineethanesulfonic acid (50 mM), sodium chloride (150 mM), Triton X-100 (1%), sodium deoxycholate (1%), sodium dodecyl sulfate (SDS;0.1%), glycerol (10%), and protease inhibitors (Complete Protease Inhibitor Cocktail, Roche, Mannheim, Germany). Lysates were cleared by centrifugation, and supernatants were boiled in SDS loading buffer. Samples were separated by SDS polyacrylamide gel electrophoresis (PAGE) and immunoblotted with indicated antibodies.

### Immunoprecipitation

Cells were lysed that includes Triton X-100 (1%), HEPES (50 mM), NaCl (150 mM), glycerol (10%), ethylene glycol tetra-acetic acid (1 mM), sodium pyrophosphate (10 mM), sodium fluoride (100 mM), sodium orthovanadate (1 mM), and protease inhibitors. Lysates were cleared by centrifugation, and equal amounts of proteins were subjected to immunoprecipitation with indicated antibodies. The lysates were rotated overnight at 4°C, then protein G or A agarose beads were added, and the mixture was rotated for 2 h at 4°C. The immunocomplexes were washed tree times with immunoprecipitation buffer and suspended with 2X SDS-loading buffer. Samples were separated by SDS-PAGE and immunoblotted with indicated antibodies.

### Genomic DNA Dot blot assay

Genomic DNA (gDNA) was extracted from breast cancer cells using QIAamp DNA mini kits according to the manufacturer’s instructions (Qiagen). gDNA samples were diluted with 2 N NaOH and 10 mM Tris-HCl, pH 8.5, and then loaded on Amersham Protran Nitrocellulose membranes 0.2 µm (GE Healthcare) using a 96-well dot-blot apparatus (Bio-Rad). After being dried at room temperature (RT) for 1 h and gDNA was crosslinked to the membrane using GS Gene Linker (Bio-Rad, Hercules, CA), then blocked with 5% nonfat milk for 1 h at RT. Membranes were incubated in primary antibodies against either 5-mC or 5-hmC at 4 °C overnight. 5-mC or 5-hmC were visualized by using chemiluminescence. To ensure equal loading, membranes were stained with methylene blue (0.02%) after immunoblotting.

### RNA extraction and quantitative real-time quantitative polymerase chain reaction

Total RNAs were isolated using a kit (RNeasy kit, Qiagen) and converted to cDNA using random hexamers and reverse transcription (Superscript III, Life Technologies, Carlsbad, CA). real-time quantitative polymerase chain reaction (RT-qPCR) was performed (CFX Connect and iTaq Universal SYBR Green SMX, Bio-Rad). All PCR reactions were performed with primers (Supplemental Table 1) with the following steps: initial denaturation, 95°C for 10 minutes; 40 cycles of denaturation, 95°C for 15 seconds and annealing/extension, 60°C for 60 seconds.

### RNA sequencing and analysis

RNA from breast cancer cells were isolated using the RNeasy kit (Qiagen). RNA was analyzed for integrity, quality and quantity using the Agilent BioAnalyzer 2100.

Three independent RNA samples per experimental groups were prepared. cDNA Library preparation and RNA sequencing were performed on an IlluminaHiSeq platform as paired-end 150-bp reads by Genewiz (South Plainfield, NJ).

Raw data quality was evaluated with FastQC, reads were trimmed using Trimmomatric v.0.36 to remove adapter sequences and poor-quality nucleotides. Reads were then mapped onto homo sapien GRCh38 reference genome using STAR aligner v.2.5.2b. Genome was available on ENSEMBL. Generated BAM files were used to extract unique gene hit counts using FeatureCounts from Subread package v.1.5.2.

Genes were identified on mapped reads followed by downstream differential expression analysis using R package DESeq2. Genes with less than 5 reads per sample were removed. Wald test was used to statistically identify genes with P value <0.05. Using regularized logarithm (rlog) function, count data was transformed for visualization on a log2 scale. Data were analyzed using Gene Set Enrichment Analysis (GSEA).

### Generation of plasmids lentivirus production

Scramble and DNMT3A shRNA were cloned in pSicoR-mCherry (#21907, Addgene)^11^. Lentivirus was made using a third-generation packaging system in HEK 293FT cells. At 72 hours after transfection, lentivirus-containing medium was centrifuged to remove cell debris and passed through a 0.45-μm filter. Lentivirus was concentrated by ultracentrifugation at 25,000 rpm for 1.5 hours using SW40-Ti rotor (Beckman Coulter, Brea, CA). Supernatant was removed and lentivirus was resuspended in sterile PBS. Lentivirus was divided into aliquots and stored at −80°C until used.

### Immunofluorescence staining

Frozen sections were fixed with cold acetone for 15 minutes and washed thrice with PBS. Samples were permeabilized with 0.1% Triton X-100 for 10 minutes, washed with PBS, and incubated with blocking solution containing 3% bovine serum albumin (BSA) and 1% goat serum for 1 hour at RT. Sections were incubated with anti-FAK (1:200), anti-pY397 FAK (1:100), anti-DNMT3A (1:100), anti-DNMT3B (1:100) overnight at 4°C. For staining of 5-mC and 5-hmC, DNA was denatured with 2N HCL for 30 min, and then neutralized in Tris-HCl (PH 7.4) for 10 min before blocking. Sections were washed with PBS and incubated with conjugated goat anti-rabbit or mouse secondary antibodies (1:1000) (Alexa Fluor 594 or 488, Thermo Fisher) for 1 hour at RT. Slides were mounted (Fluoromount-G, SouthernBiotech, Birmingham, AL), and images were acquired with a confocal microscope at 60-fold magnification (Nikon A1R, Nikon, Tokyo, Japan).

### Statistical analysis

Data sets underwent Shapiro-Wilk test for normality, and statistical significance between experimental groups was determined with student *t*-test or two-way analysis of variance (2-way ANOVA) with Sidak multiple comparisons test (Prism software, v7.0d; GraphPad Software, La Jolla, CA). Power analyses were performed to determine sample size for 2-way ANOVA.

## RESULTS

### FAK inhibition reduces breast cancer cell proliferation and DNA methylation

To elucidate the biological role of FAK activation in breast cancer proliferation, we performed cell proliferation assay using MCF7 and MDA231 breast cancer cells treated with FAK inhibitor (FAK-I). FAK-I significantly reduced MCF7 and MDA231 cell number (S. Fig. 1A). We further evaluated FAK-I effect on cell proliferation by staining for the proliferation marker Ki67.^12^ Consistent with these results, we observed a dramatic decrease in the mean number of Ki67 positive cells/field after 24 h treatment of FAK-I compared to vehicle treated (Figure 1A).

**Figure 1.**
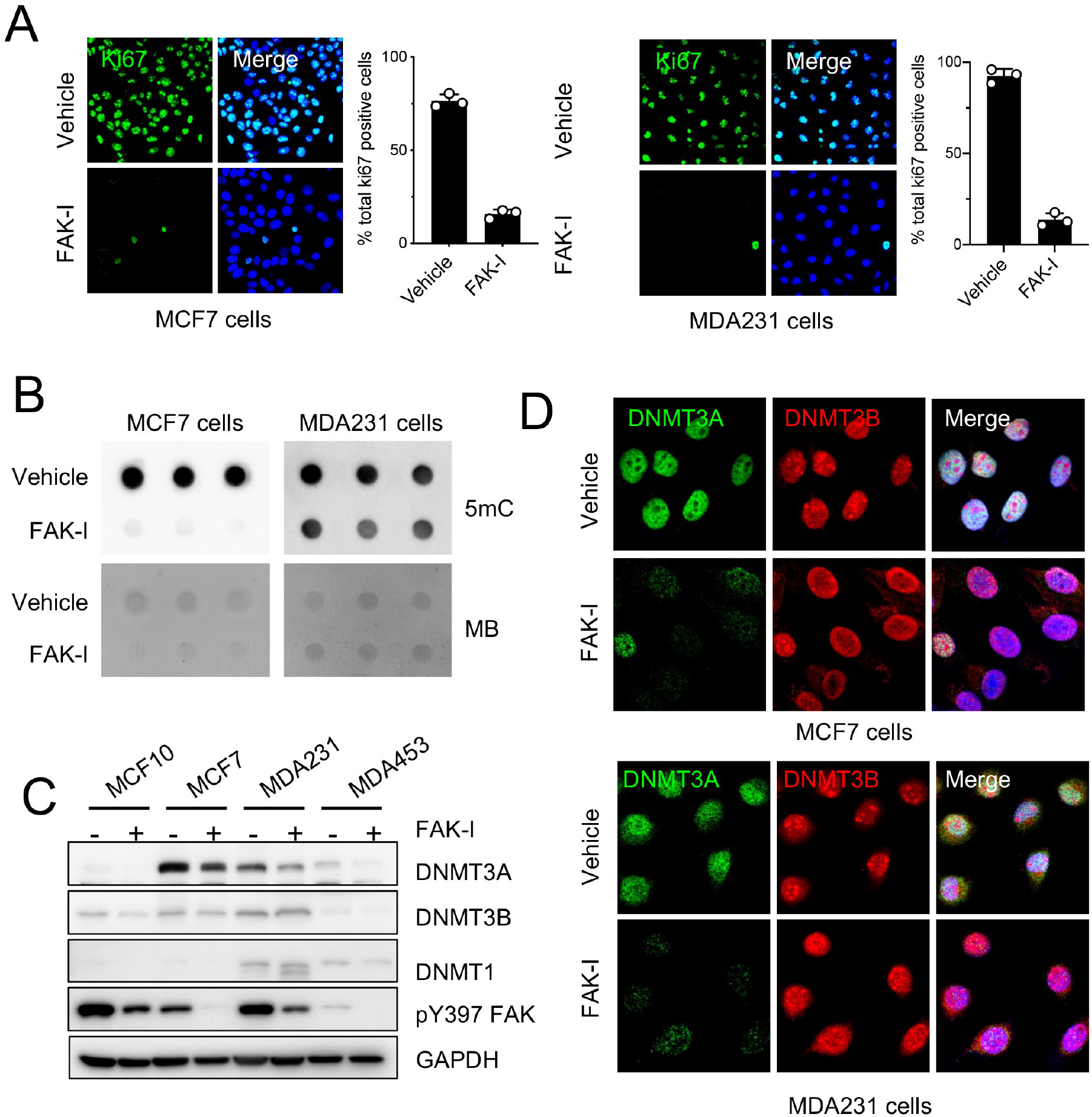
Loss of FAK activity reduces breast cancer cell proliferation and global DNA methylation via reducing DNMT3A. **A**, MCF7 and MDA231 cells were treated FAK-I (VS-6063, 2.5 μM) for 2 days. Representative immunostaining of Ki-67 (green) and blue (DAPI) were merged. Scale bars, 20 μm. The percentage of Ki-67 positive cells was calculated for each square separately (± SEM, n=3, ^**^P<0.005 vs. vehicle, unpaired t-test). **B**, Dot blots for 5-mC using genomic DNA (gDNA) from MCF7 and MDA231 treated with/without FAK-I (2.5 μM) for 2 days. Methylene blue staining of 150 ng total gDNA was used to loading control (n=3). **C**, Breast cancer cells were treated with FAK-I (VS-6063, 2.5 μM) for 24 h. Shown are immunoblots of lysates for DNMT3A, DNMT3B, DNMT1, pY397 FAK, and GAPDH as loading control (n=3). **D**, MCF7 and MDA231 cells were treated with FAK-I (2.5 μM) for 24 h. Immunofluorescence staining for DNMT3A (green), DNMT3B (red), and DAPI (blue) are shown (n=3).

One of the major drivers in cancer cell proliferation can be attributed to DNA hypermethylation, an epigenetic modification that leads to silencing of tumor suppressor genes.^13,14^ In an effort to test whether FAK-I might change DNA methylation in breast cancer cells, we used a DNA dot-blot assay to measure global 5-methylcytosine (5-mC) in breast cancer cell lines. Dot blotting of MCF7 and MDA231 genomic DNA for 5-mC demonstrated that pharmacological FAK-I decreased 5-mC levels in both MCF and MDA231 cells (Figure 1B). As 5-mC is catalyzed by DNA methyltransferase (DNMTs), we first investigated if FAK inhibition reduced expression of DNMTs. Of the DNMTs known to methylate cytosine at CpG sites, FAK-I specifically reduced DNMT3A but not DNMT1 or DNMT3B expression in several human breast cancer cell lines (Figure 1C). Further, FAK-I decreased pY397 FAK staining and increased FAK nuclear localization in MC7 and MDA231 cells (S. Fig. 1B,C). This was correlated with a decrease in DNMT3A staining (Figure 1D), suggesting that FAK-I-induced nuclear localization reduces DNMT3A expression.

### FAK-I promotes DNMT3A proteasomal degradation in breast cancer cells

Like human breast cancer cells, we found that FAK-I can reduce Dnmt3a levels in the 4T1 murine breast cancer cell line (Figure 2A). However, despite observing decreased Dnmt3a protein expression, FAK-I did not alter expression of *Dnmt3a, Dnmt3b*, or *Dnmt1* mRNA expression (Figure 2B), suggesting FAK-I alters Dnmt3A protein stability. To confirm the possibility whether FAK regulates Dnmt3a posttranscriptionally, 4T1 cells were treated with combinations of FAK-I and the proteasomal inhibitor MG132. While FAK-I alone reduced DNMT3A levels, co-treatment with FAK-I and MG132 failed to reduce DNMT3A (Figure 2C). To decipher how FAK regulates Dnmt3a, but not Dnmt3b or Dnt1 expression, we performed FAK-Dnmt interaction study in 4T1 cells. Endogenous FAK-Dnmt3a interaction in 4T1 cells was confirmed from FAK immunoprecipitation (IP) following FAK-I treatment (Figure 2D).

**Figure 2.**
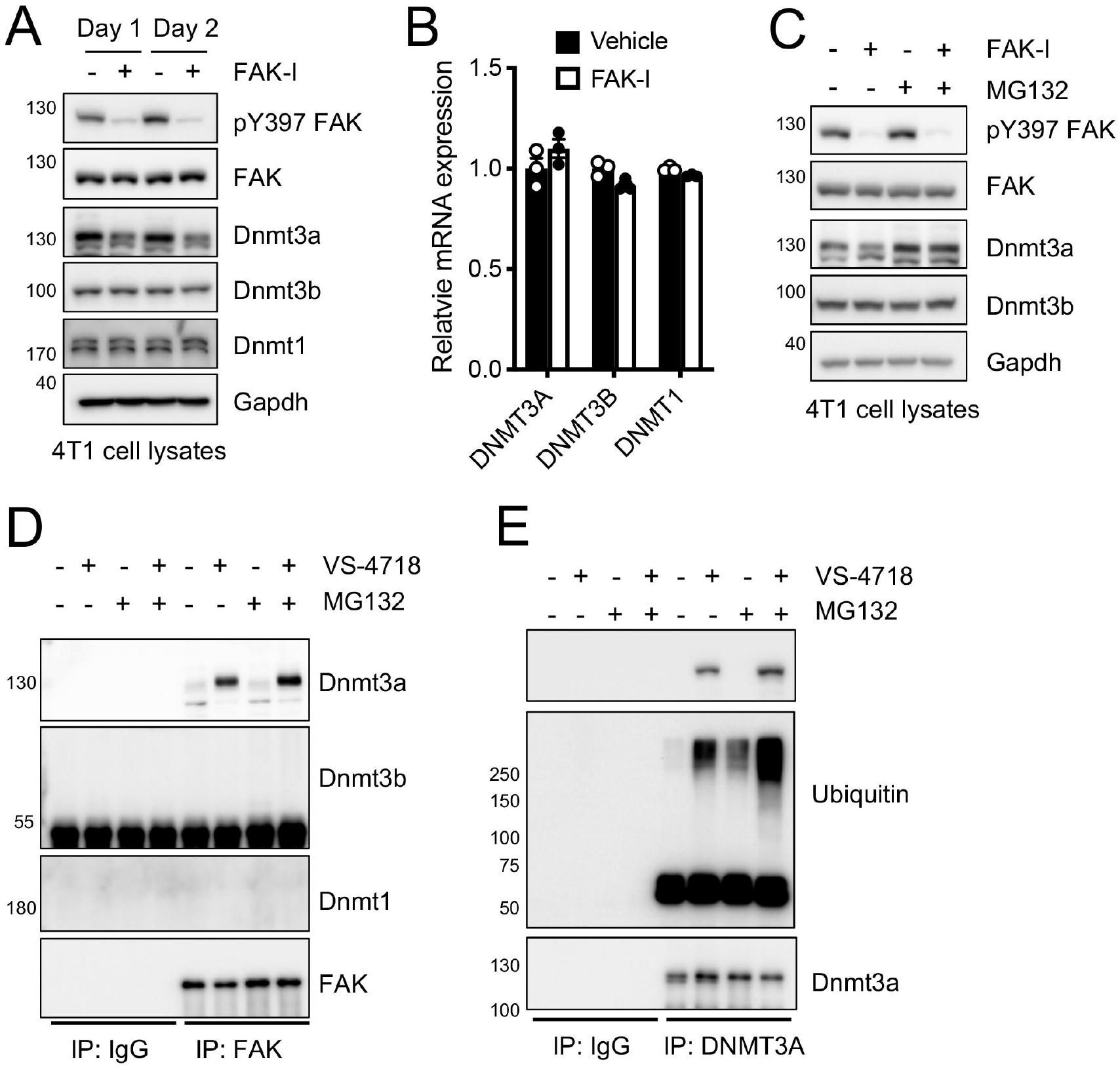
FAK-I increases proteasomal degradation of Dnmt3a in breast cancer cells. **A**, 4T1 cells treated with FAK-I (2.5 μM) for indicated time. Shown are immunoblots of lysates for pY397 FAK, FAK, Dnmt3a, Dnmt3b, Dnmt1, and Gapdh as loading control (n=3). **B**, mRNA levels of *Dnmnt*s were measured after 1 day FAK-I (2.5 μM) treatment (±SEM, n=3). **C-E**, 4T1 cells were treated with FAK-I (2.5 μM) only or together with MG132 (20 μM) for 12 h. Whole cell lysates were immunoprecipitated with anti-FAK or anti-DNMT3A antibody and subjected to immunoblotting with indicated antibodies (n=3).

Association of FAK and Dnmt3a was further increased when 4T1 cells were treated with the proteasomal inhibitor MG132 in addition to FAK-I (Figure 2D). However, we failed to detect Dnmt3b and Dnmt1 in FAK IP (Figure 2D). Anti-ubiquitin immunoblotting of Dnmt3a IP in 4T1 cells showed that FAK-I increased ubiquitination of Dnmt3a, which was further enhanced by MG132 co-treatment (Figure 2E). These results demonstrate that FAK inhibition promotes DNMT3A ubiquitination and proteasomal degradation in breast cancer cells.

### FAK inhibition promotes tumor suppress gene DAB2 expression

To determine if DNMT3A expression is important for breast cancer cell proliferation, we designed short hairpin RNA (shRNA) to knockdown DNMT3A in MCF7 and MB231 cells. Knockdown of DNMT3A was evaluated at both the protein and mRNA level (S. Fig. 2A,B). There was no change in DNMT3B or DNMT1 expression (S. Fig. 2A). Proliferation assay of shScramble (shScr) and shDNMT3A cells showed that loss of DNMT3A could reduce cell growth (S. Fig. 2C). To better understand how FAK-I or DNMT3A knockdown could reduce breast cancer cell proliferation we performed RNA sequencing on vehicle, FAK-I, shScr, and shDNMT3A MCF7 cells. We found that 386 and 604 genes were upregulated in FAK-I and shDNMT3A MCF7 cells compared to their controls, respectively (Figure 3A). Of these upregulated genes, 138 were found to be upregulated in both conditions (Figure 3B). We analyzed potential pathways altered by these upregulated genes using Gene Set Enrichment Analysis (GSEA). GSEA revealed the enrichment of common genes were associated with HALLMARK TNFA SIGNALING VIA NFKB, HALLMARK KRAS SIGNALING UP, HALLMARK IL2 STAT5 SIGNALING, and HALLMARK TGF BETA SIGNALING (Figure 3C). As transforming growth factor-β (TGF-β) pathway is an important regulator of breast cancer progression,^15^ we further analyzed this data set. Heat maps of genes with significant changes in the TGFBR_PATHWAY that were affected by FAK-I and shDNMT3A knockdown (Figure 3D). We decided to focus on Disabled-2 (*DAB2*) which was identified as upregulated in both FAK-I and shDNMT3A MCF7 cells (Figure 3D). DAB2 is a tumor suppressor that has been shown to be down regulated in several types of cancer.^16-19^ We compared *DAB2* expression in normal and breast invasive carcinoma (BRCA) samples using the GEPIA (Gene Expression Profiling Interactive Analysis) dataset (http://gepia.cancer-pku.cn/). The results indicated that the expression levels of *DAB2* was lower in BRCA than in paired normal samples (Figures 3E). Next, we tested whether FAK activity regulate DAB2 expression in breast cancer cell lines. Analysis of mRNA levels revealed that FAK-I promoted *DAB2* expression through increased transcription (Figure 3F). DAB2 expression was further verified at the protein level following FAK-I treatment of different breast cancer cell lines (Figure 3G and S. Fig. 3A). Knockdown of DNMT3A using shRNA also increased DAB2 protein expression (S. Fig. 2A and D), suggesting that FAK-I increases DAB2 via decreased DNMT3A stability. It is well-known that there is a negative correlation between DNA methylation and gene expression and the CpG island of DAB2 promoter has been reported to be methylated in various cancers.^16,20-23^ According to the MethHC database (http://methhc.mbc.nctu.edu.tw/php), the DAB2 promotor is hypermethylated in BRCA compared to normal samples, thereby leading to the downregulation or silencing of DAB2 (Figure 3H). Since FAK-I increased *DAB2* at a transcriptional level (Figure 3F), we next investigated if FAK-I treatment decreased 5-mC levels within the promoter of *DAB2*. 5-mC immunoprecipitation experiments revealed that FAK-I treatment reduced 5-mC on the DAB2 promoter in breast cancer cells by increasing the non-methylated DNA fraction (S. Fig. 3B). These data suggest that FAK inhibition promotes increased DAB2 expression through decreased promoter methylation in breast cancer cells.

**Figure 3.**
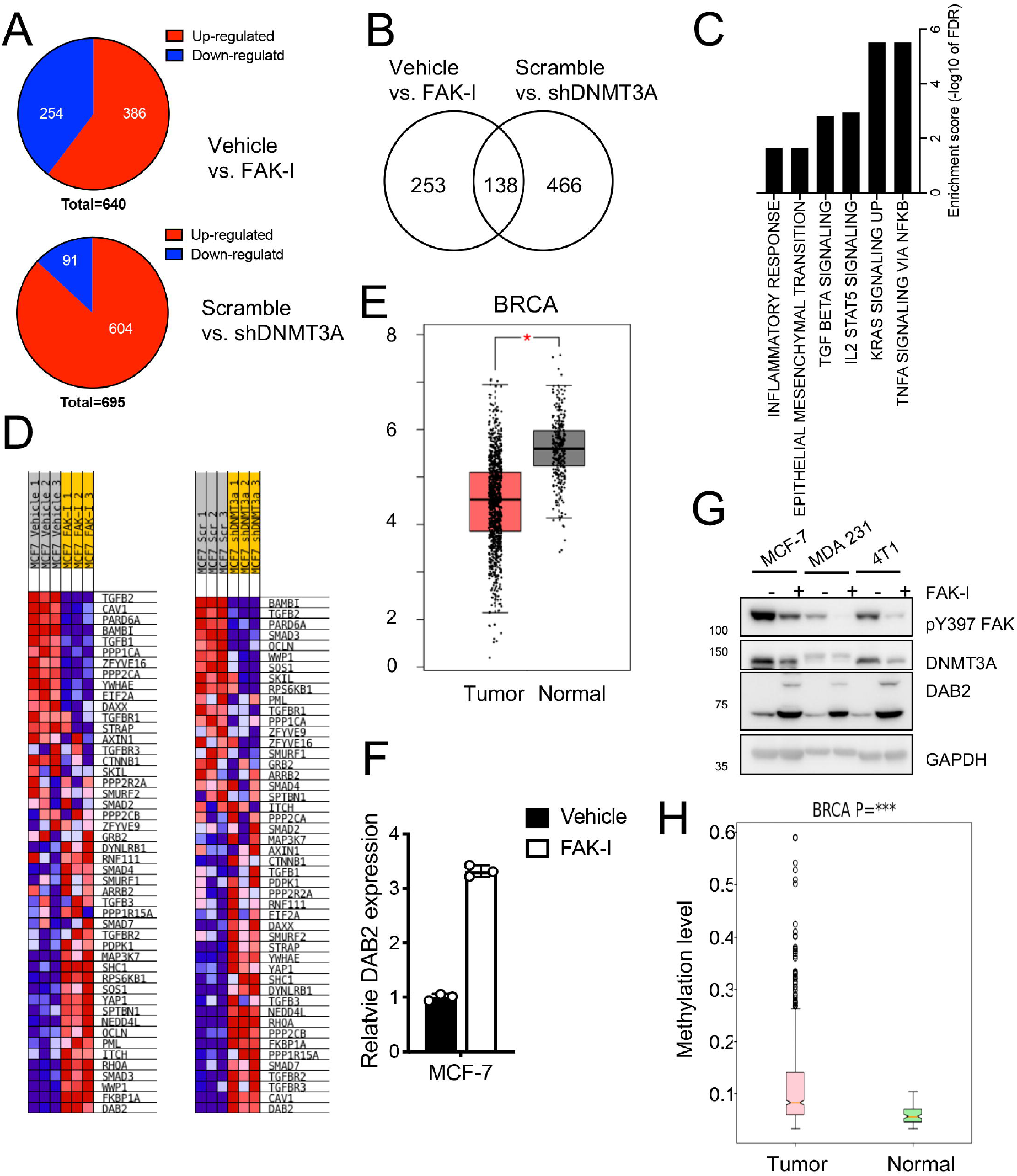
DNMT3A-regulated gene expression signatures in breast cancer cells. MCF7 cells were treated with FAK-I (VS-6063, 2.5 μM) for 2 days or infected DNMT3A shRNA (shDNMT3A) lentivirus. RNA sequencing data were analyzed. **A**, Pie chart depicting the number of FAK activity or DNMT3A-regulated genes in MCF7 cells. **B**, Venn diagram indicating overlap up-regulated genes identified between FAK inhibition and shDNMT3A. **C**, Comparison of gene set expression scores in common up-regulated genes between FAK-I treated and shDNMT3A MCF7 cells. Gene sets include the MSigDB hallmark collection. **D**, Heatmap of the genes contributing to TGFR pathway GSEA enrichment plot. **E**, mRNA expression levels of DAB2 was analyzed by GEPIA platform. DAB2 were confirmed significantly higher in normal samples than that in breast cancer samples. ^*^ *P*<0.05 vs. tumor. **F**, mRNA levels of DAB2 were measured after FAK-I (VS-6063, 2.5 μM) treatment for 2 days in MCF7 cells (±SEM, n=3, unpaired t-test, ^**^ *P*<0.005). **H**, Comparison DNA methylation of DAB2 gene promoter in breast cancer samples and matched normal samples. Data were taken from the MethHC database (http://methhc.mbc.nctu.edu.tw/). ^*^*P*< 0.0001.

**Figure 4.** FAK inhibition reduces tumor growth and DNMT3A leading to upregulation of DAB2. 4T1 cells were implanted into the mammary fat pad of BALB/c mice. Mice were treated with vehicle or FAK-I (VS-4718, 50 mg/kg) twice daily via oral gavage starting on day 12. Tumor formation and its growth were monitored. **A**, Volume of tumors were measured (±SD, n=5, unpaired t-test vs. vehicle, ^*^ *P*<0.05, ^**^ *P*<0.01). **B**, Shown are representative images of tumor collected at day 20. **C**, Weight of tumors were measured (±SD, n=5, unpaired t-test vs. vehicle, ^*^ *P*<0.05, ^**^ *P*<0.01). **D**, mRNA levels of *Dab2* were measured (±SEM, n=3). **E**, Shown are immunofluorescence staining of tumor sections for FAK, pY397 FAK, Dnmt3a, Dnmt3b, 5-mC, 5-hmC, and Dab2 (n=4). Red, green, and blue (DAPI) were merged. Scale bars: 100 μm.

### FAK inhibition reduces 4T1 breast cancer growth in vivo

To evaluate if FAK inhibition could reduce breast cancer cell proliferation in vivo, we injected 4T1 breast cancer cells into the mammary fat pads of mice and, 12 days after injection, started dosing the mice with vehicle or FAK-I. Mice treated with FAK-I displayed reduced tumor growth by approximately 50% as early as day 16 compared to the vehicle-treated tumors compared to vehicle treated mice (Figure 5A). Additionally, final tumor size and weight were significantly decreased in FAK-I treated mice compared to vehicle (Figure 5A-C). *Dab2* mRNA expression within 4T1 tumors was analyzed and found to be elevated in the FAK-I treated group compared to vehicle (Figure 5D). To determine if FAK-I altered Dnmt3a, 5mc, 5hmC, and Dab2 expression, we performed immunostaining via 4T1 tumors. While vehicle treated tumors showed primarily cytoplasmic FAK localization, treatment with FAK-I increased FAK nuclear localization within tumors (Figure 5E). Further, high levels of active pY397 FAK were observed in vehicle treated tumors compared to low levels in the FAK-I group (Figure 5E). Increased FAK nuclear localization in tumors was associated with decreased Dnmt3a expression, but not Dnmt3b and Dnmt1 (Figure 5E). The results showed that levels of 5mC was decreased and negatively correlated with 5hmC and Dab2 by FAK inhibition, supporting our findings that DAB2 are transcriptionally repressed by DNMT3A.

**Figure 5.**
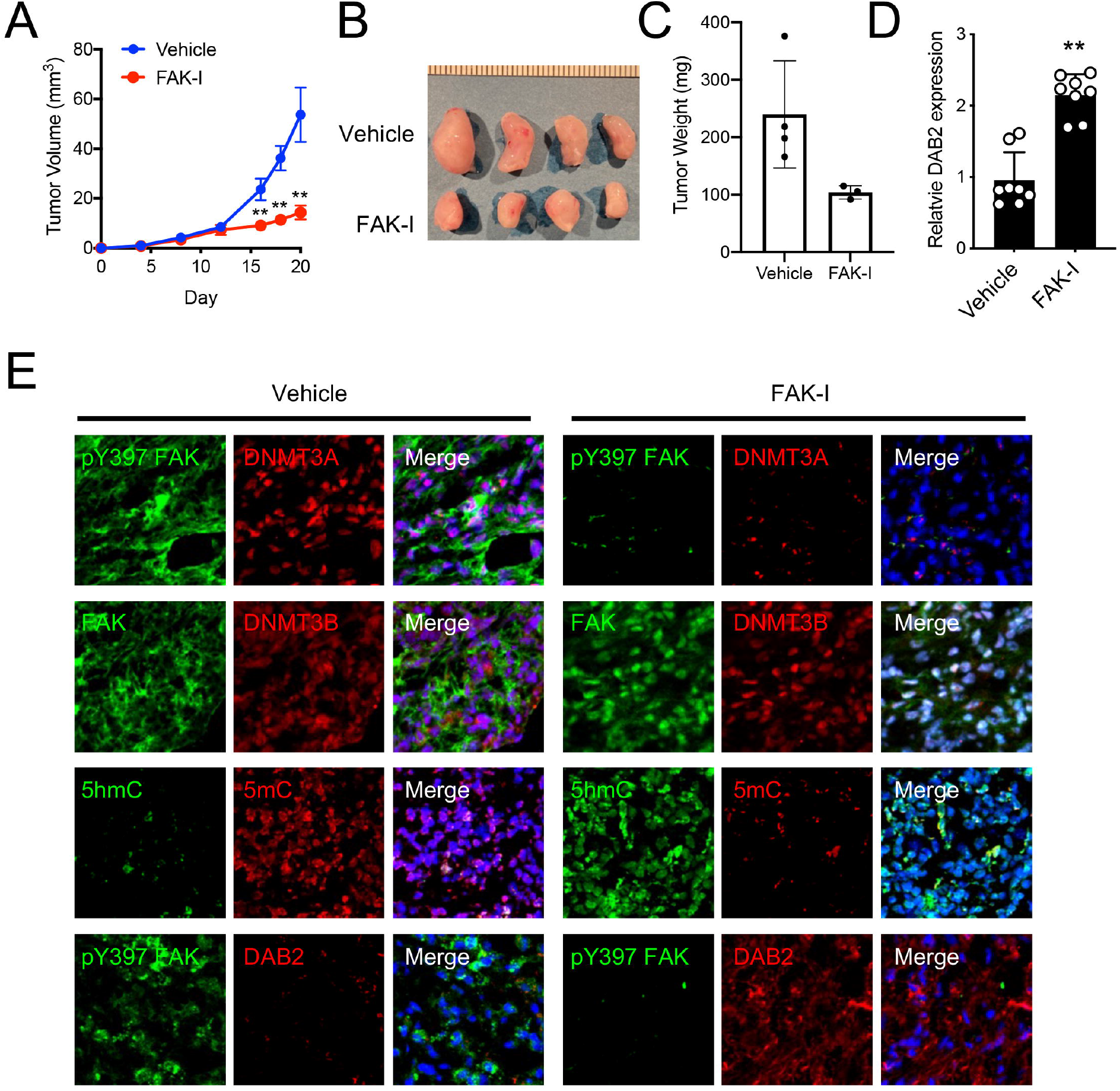

## Discussion

In this study, we found that FAK inhibitor (FAK-I)-induced nuclear FAK promotes DNMT3A degradation via ubiquitination and altering TGFβ receptor (TGFβR) signaling to reduce cancer cell survival. We focused on increased DAB2 gene expression by treating human and mouse breast cancer lines with FAK-I. As other cancer cells also express both DNMT3A and FAK, we expect that other human tumors would also upregulate DAB2 following FAK-I treatment which can be used as an anti-tumor agent.

While the molecular process behind FAK nuclear localization is unknown, loss of FAK activity through pharmacological or genetic inhibition or loss of integrin-mediated adhesion leads to elevated nuclear FAK localization. In the nucleus, we found that FAK can interact with DNMT3A, but not DNMT3B or DNMT1, which is consistent with our previous work in vascular smooth muscle cells.^24^ This is interesting as DNMT3A and DNMT3B and structurally similar and drive de novo methylation, suggesting that DNMT3A and DNMT3B may not have overlapping or compensatory roles in the gene promoters they silence. Additionally, DNMT1 mediates DNA fingerprinting by copying DNA methylation states onto newly synthesized strands,^25^ and will not remethylate DAB2 promoters under FAK-I.

TGFβ signing in cancer can either increase or decrease cancer cell survival.^26^ DAB2 has been shown to act as a molecular switch to alter TGFβ signaling.^27^ In the absence of DAB2, TGFβ activation of TGFβR leads to cell proliferation, migration, and survival.^27^ Mechanistically in the noncanonical pathway, activation of TGFβR recruits Shc and Grb2. Without DAB2, Grb2 recruits SOS which leads to Ras/MAPK signaling to promote Cell survival and proliferation.

DAB2 expression binds Grb2, preventing SOS recruitment, which promotes canonical TGFβR-SMAD pathway to promote cell death and growth arrest. However, it remains to be seen if increased DAB2 expression following FAK-I treatment is the driver of reduced cell proliferation.

While FAK inhibition may influence many signaling pathways or gene expression, we found lentiviral mediated knockdown of DNMT3A via shRNA could also increase DAB2 expression and inhibit cell proliferation. These data suggest that DNA methylation mediated by DNMT3A is what suppresses DAB2 in breast cancer. Therefore, DNMT inhibitors may also be used to induce DAB2 expression and reduce tumor cell proliferation.

## Abbreviations

5-mC: 5-methylcytosine
DNMT: DNA methyltransferase
FAK: focal adhesion kinase
FAK-I: FAK inhibitor
DAB2: disabled homolog 2
gDNA: genomic DNA
pY397: autophosphorylation at tyrosine 397
RT-qPCR: real-time quantitative polymerized chain reaction
shRNA: short hairpin RNA

## Author Contributions

JMM and KJ, performed *in vitro* assays. JMM and KJ performed the animal studies and analysis. JMM, KJ, EYA, and STL provided data interpretation and analysis. JMM, KJ, and STL wrote the manuscript with input from all authors.

