## Supplementary figures and images for "FAK inhibition suppresses breast cancer progression via DNA methylation-mediated DAB2 gene reactivation"

### Supplemental Figure 1

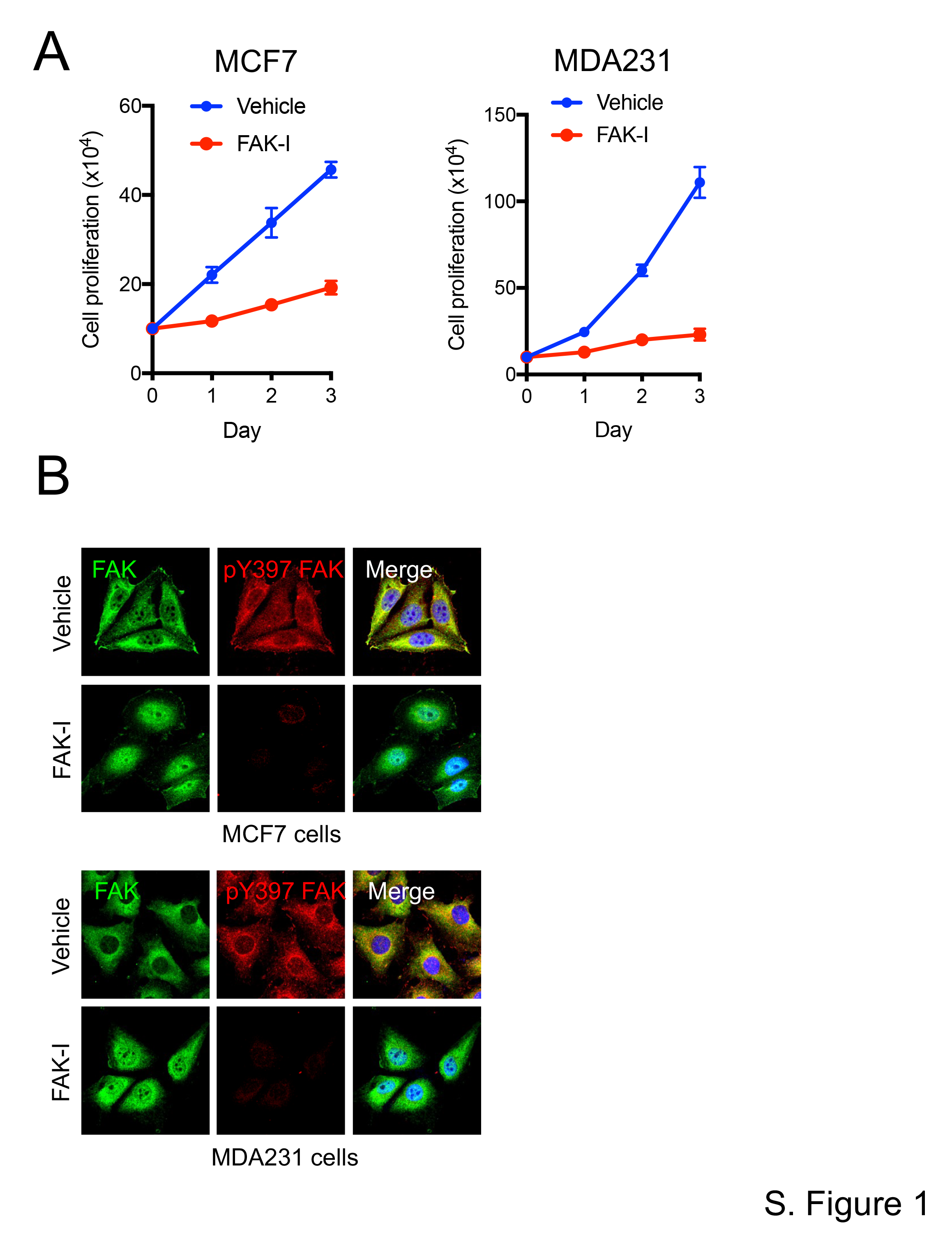

### Supplemental Figure 2

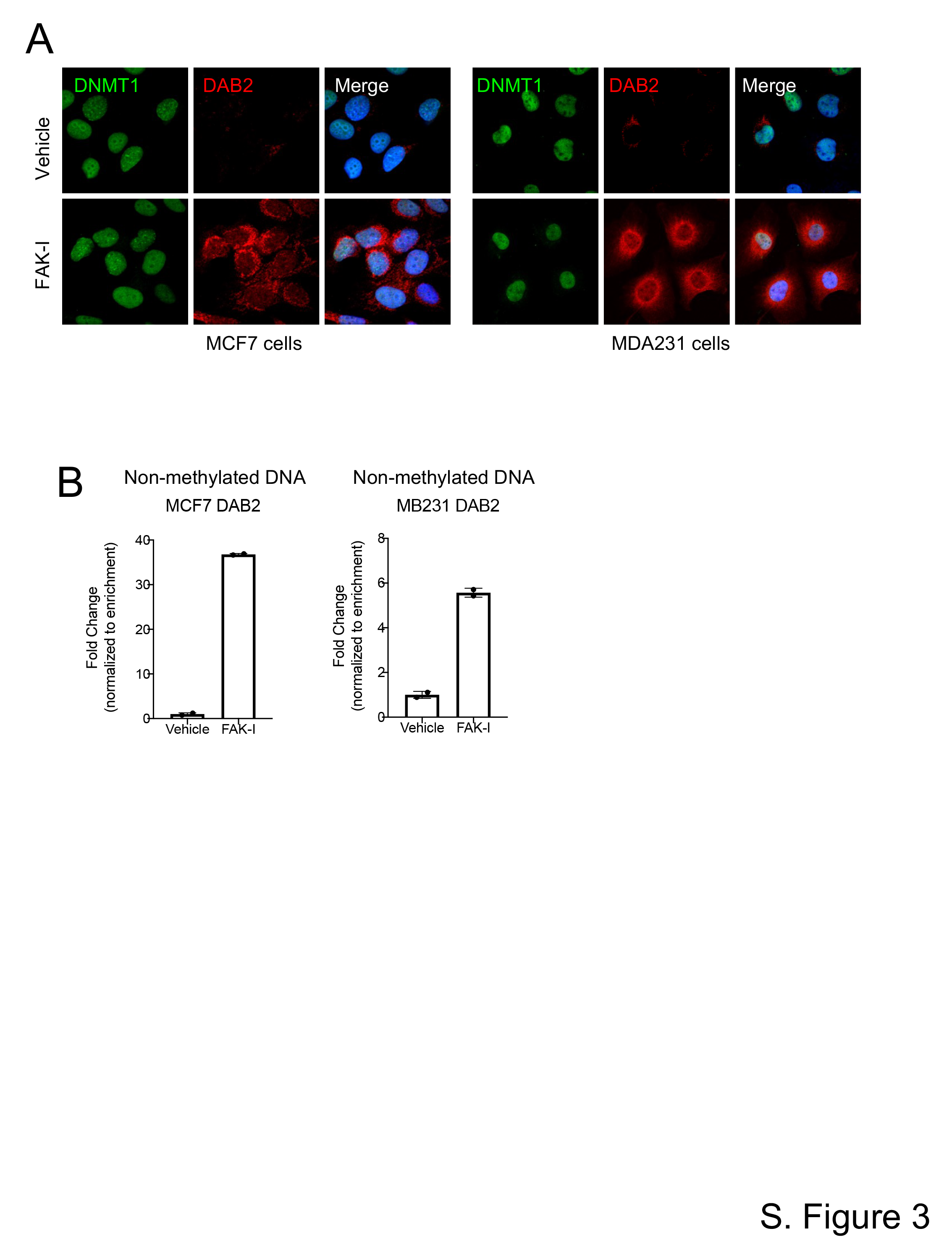

### Supplemental Figure 3

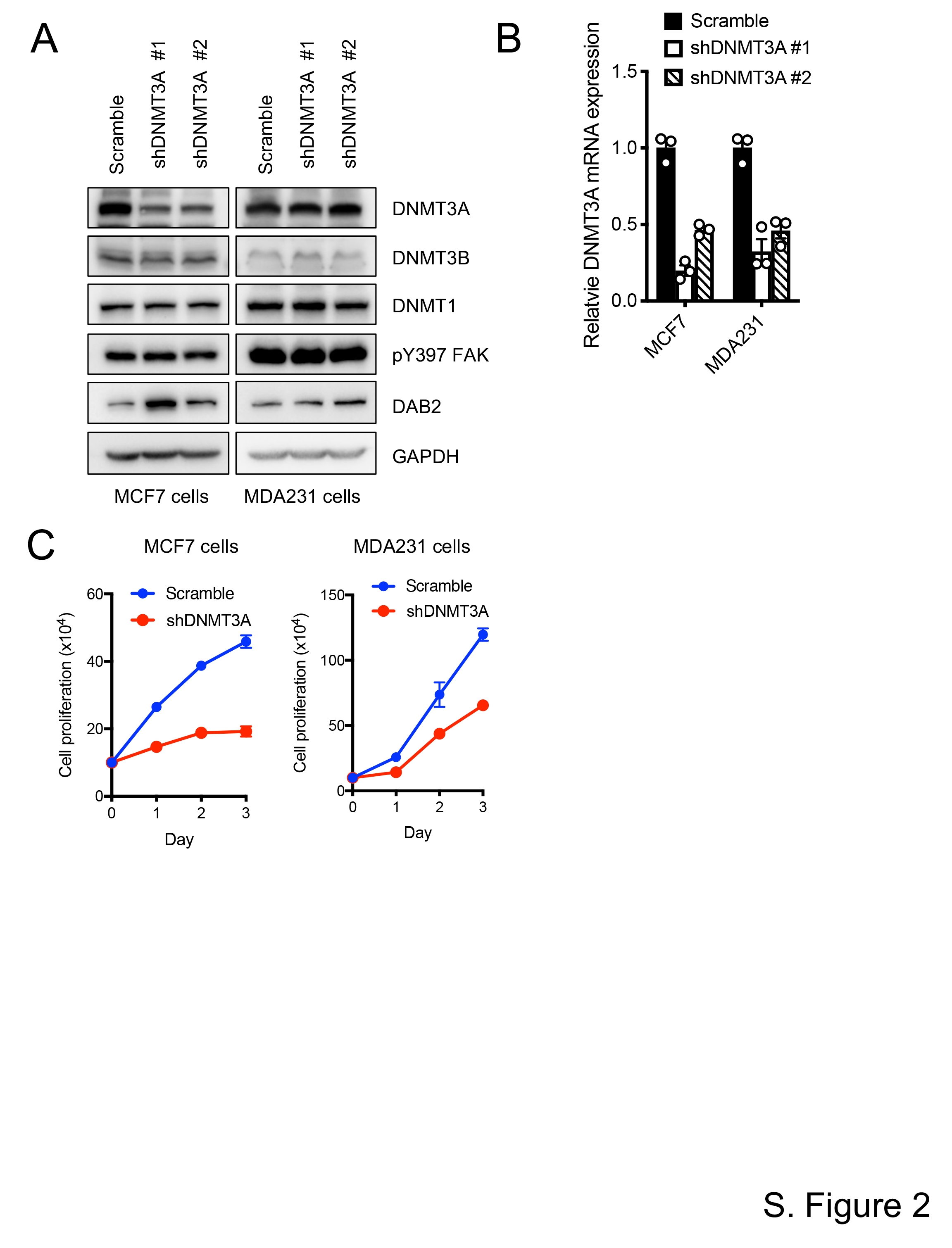
