## Supplemental Figure Legends for "FAK inhibition suppresses breast cancer progression via DNA methylation-mediated DAB2 gene reactivation"

**S. Figure 1. FAK inhibition blocks the proliferation of breast cancer cells. A,** MCF7 and MDA231 cells were treated FAK-I (VS-6063, 2.5 μM) and cells were counted on the indicated days (± SD, n=3, ***P*<0.005 vs. Vehicle, two-way ANOVA followed by Sidak multiple comparisons test). **B,** Immunostaining of FAK and pY397 FAK were performed with/without FAK inhibitor (VS-6063, 2.5 μM) for 24 h. Red, green, and blue (DAPI) were merged. Scale bars: 20 μm.

**S. Figure 2. Knock-down of DNMT3A in breast cancer cells reduces proliferation via inducing DAB2.** MCF7 and MDA231 cells were infected with shDNMT3A lentivirus. **A,** Shown are immunoblots of lysates for DNMT3A, DNMT3B, DNMT1, pY397 FAK, DAB2, and GAPDH as loading control (n=3). **B,** mRNA levels of DNMT3A were measured (±SEM, n=3). **C,** Cells were counted on the indicated days (± SD, n=3, ** *P*<0.005 vs. Vehicle, two-way ANOVA followed by Sidak multiple comparisons test).

**S. Figure 3. FAK-I increases DAB2 expression and reduces *DAB2* promoter methylation. A,** MCF7 and MDA231 cells were treated FAK-I (VS-6063, 2.5 μM) for 24 h. Immunostaining for DNMT1 (green), DAB2 (red), and nuclei (DAPI, blue) (n=3). **B**, Methyl Collector assay showing reduced methylation at the GADD45A promoter in FAK-I (2.5 μM, 2 days) treated MCF7 and MDA231 cells (±SEM, n=3, unpaired t-test, ** *P*<0.005)
